# A framework for collaborative computational research

**DOI:** 10.1101/033654

**Authors:** Apuã C. M. Paquola, Jennifer A. Erwin, Fred H. Gage

## Abstract

Analysis of high troughput biological data often involves the use of many software packages and in-house written code. For each analysis step there are multiple options of software tools available, each with its own capabilities, limitations and assumptions on the input and output data. The development of bioinformatics pipelines involves a great deal of experimentation with different tools and parameters, considering how each would fit to the big picture and the practical implications of their use. Organizing data analysis could prove challenging. In this work we present a set of methods and tools that aim to enable the user to experiment extensively, while keeping analyses reproducible and organized. We present a framework based on simple principles that allow data analyses to be structured in a way that emphasizes reproducibility, organization and clarity, while being simple and intuitive so that adding and modifying analysis steps can be done naturally with little extra effort. The framework suppports version control of code, documentation and data, enabling collaboration between users.

## INTRODUCTION

Computational analyses of biological data, especially those involving high-throughput techniques such as next-generation sequencing, often require the use of multiple software tools and in-house written code. For each step of an analysis, there are often multiple options of software tools available, each tool having its own capabilities, limitations and assumptions on the structure of input and output files. The development of bioinformatics pipelines involves a great deal of experimentation with different tools and parameters, to test how each would fit the big picture, considering what each would add to the analysis and the practical implications of their use. Due to the diversity of computational tools and setups it can prove challenging to keep data analysis organized and reproducible.

Several efforts have been made to document orgazination principles for computational research and develop tools to manage and automate analysis workflows [Noble et al. 2009, Ebert et al. 2015, Goodstadt L. 2010, Köster et al. 2010, Goecks et al. 2010]. A prominent example is the widely used Galaxy patform [Goecks et al. 2010], which allows the user to compose a workflow by linking inputs and outputs of tools in a graphical user interface. Galaxy addresses reproducibility by the use of well-defined wrappers for each tool and automatic recalculation of intermediate results. The drawbacks are the complexity of mantaining a local Galaxy installation, and the overhead cost of writing and modifying wrappers as new analysis tools are added.

We present a framework based on simple principles that allow data analyses to be structured in a way that emphasizes reproducibility, organization and clarity, while being simple and intuitive so that adding and modifying analysis steps can be done naturally with little extra effort. The framework suppports version control of code, documentation and data, enabling collaboration between users.

## PRINCIPLES

In this framework, the layout of data files, scripts and documentation in a file system is key to organized and reproducible data analysis. We use three principles to guide the layout of files: (1) separation of human-generated data from computer-generated data; (2) tree structure; (3) driver scripts.

A **driver script** [Noble et al. 2009] is a computer program that runs without arguments, producing an output given its input data. Running without arguments ensures the analysis doesn’t depend on manual input.

Most data analyses are composed of multiple steps that depend on each other. In a hypothetical example, steps B and C take as input the output of a previous step A. A natural way to organize these direct dependencies is to use a **tree structure** in which node A has two children nodes B and C.

The third principle is to have a clear **separation** between human-generated data and computer-generated data. Files written by humans, i.e. driver scripts, programs and documentation, specify how output files are generated, and are placed in a separate directory than computer-generated files.

Figure 1 shows a directory structure that puts these principles together, mapping the analysis steps to directories in a file system. Each node or directory contains two special subdirectories, the “human” directory “__” (double underscore), the “machine” directory “_” (single underscore), and one subdirectory for each of the children nodes. The human directory “__” contains the driver script, documentation and code. Running the driver script generates the contents of the machine directory “_”. Dependencies other than the parent are specified by putting symbolic links to other nodes in the “__/dep/” subdirectory.

**Fig. 1:**
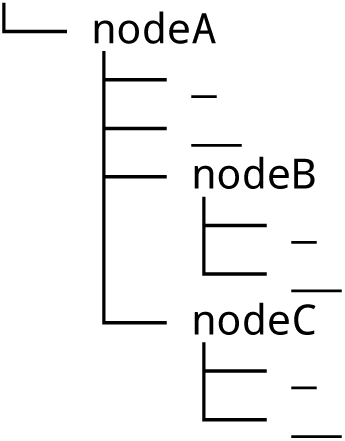
Directory structure of a hypothetical analysis tree. Each analysis step is mapped to a directory that contains two special subdirectories: __ (double underscore) for human-generated files, and _ (single underscore) for computer-generated files (program outputs). A driver script located at the “human” directory __ specifies how the contents of the “machine” directory _ are generated. The directory structure reflects data dependencies: analysis steps B and C depend on the output of step A.

With this structure, the position of a node in the tree, or equivalently the full path to a directory, documents the main analysis steps taken to generate the data in that node. In other words, the full path provides a quick and intuitive view on the analysis workflow.

Moreover, the use of a directory (“_”) for program output makes no assumption on the format of output data, which is particularly important in bioinformatics, where there’s a wide variety of tools, file formats and ways to generate output.

This structure also allows automated execution of driver scripts in the whole tree, in an order defined by topologically sorting the dependency graph. In the current implementation this is done by generating a ‘makefile’ and passing it to the unix tool ‘make’. It is not the purpose of this framework to define how to run jobs in parallel or how to farm out jobs in a distributed computer environment. This is done in conjunction with other tools, such as snakemake [Köster et al. 2012], called from the driver scripts.

## VERSION CONTROL AND COLLABORATION

Version control is an essential component in software development, as it enables experimentation with code while keeping track of the full development history. The same applies to data analysis, although version control for data is not yet as widely adopted as it is for code. In this framework we use the git [http://git-scm.com/] version control system for the “human” directories __/ a git-annex to for the machine directories _/. Git-annex [https://git-annex.branchable.com/] is an extension written for git to support version control of large files. This way we get the full functionality of git, i.e., rollback to a previous version, branching, etc., to analysis trees. One of the main features of git is to enable collaboration between users, allowing them to work each on a local copy of the code repository, and merging their changes. In this framework, users can have full local copies of the analysis tree structure and development history, i.e., the “human” directories plus the git repository, and download the contents of “machine” directories on demand, avoiding wasting storage or bandwidth. Figure 2 illustrates these ideas with a hypothetical use case.

**Figure 2:**
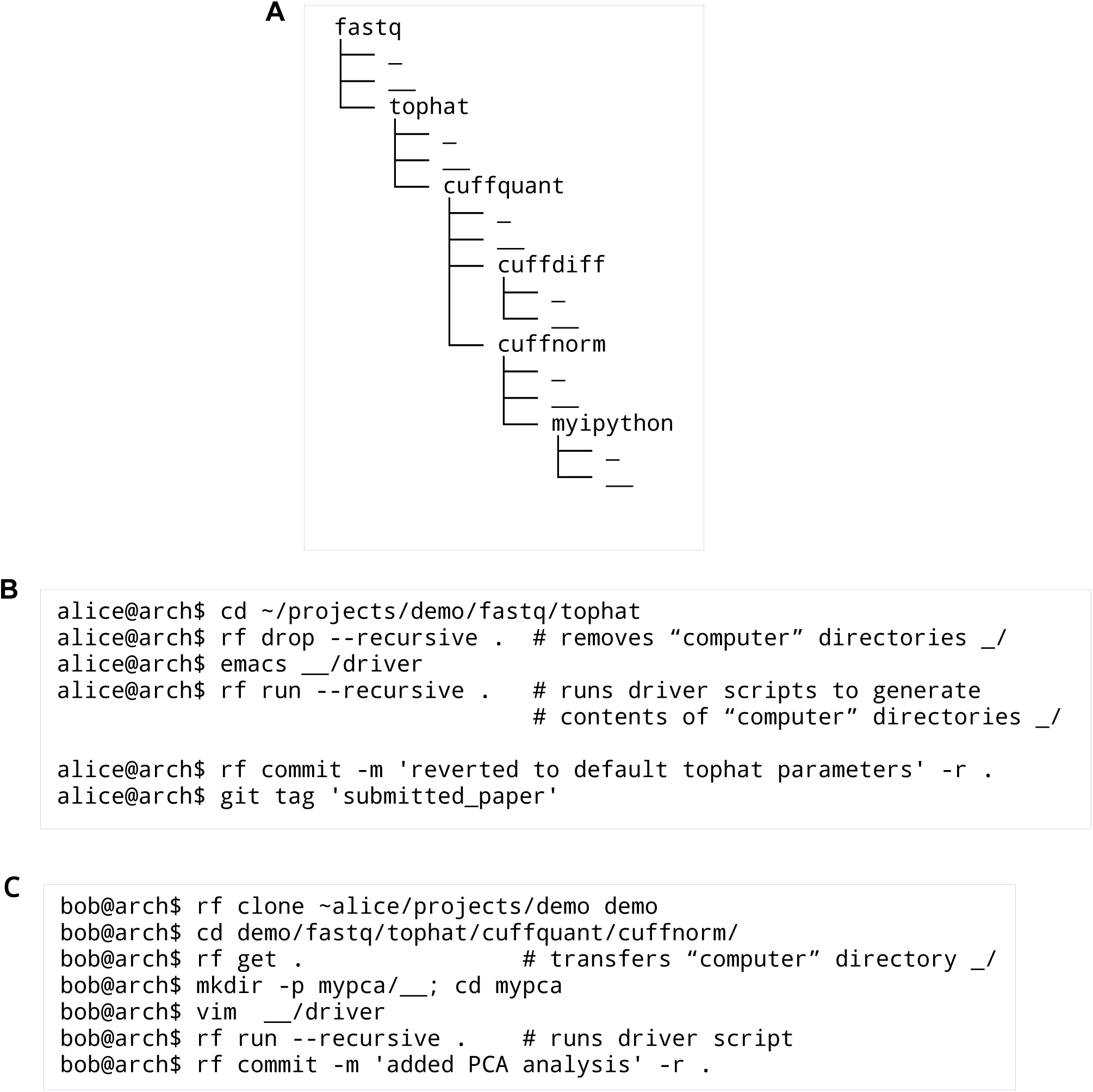
(A) Directory structure of an RNA-Seq analysis based on tophat, cuffquant and cuffnorm [Trapnell et al. 2012 and http://cole-trapnell-lab.github.io/cufflinks/manual/]. Input fastq files are located in the fastq/_directory. After execution of the driver script fastq/tophat/__/driver, the directory fastq/tophat/_contains bam files that are the output of tophat program. (B) and (C) hypothetical use cases of this framework by collaborating users Alice and Bob. The rf command is used to run driver scripts and as a front end to git and git-annex. (B) Alice edits a driver script, recomputes an analysis subtree and commits changes to version control. (C) Bob clones Alice’s analysis tree structure to a local directory, transfers data from the “cuffnorm” node, adds a new analysis node “mypca”, runs its driver script and commits changes to version control.

## PRESERVING THE EXECUTION ENVIRONMENT

In addition to the principles presented above and version control, to achive a higher degree of reproducibility, it is desirable to faithfully preserve the computational environment used in the analyses. To enable that, a feature currently in development in this framework is integration with Docker [https://www.docker.com/]. Docker provides a lightweight alternative to virtual machines to have a defined computational environment (operating system and installed software) in a software container. Docker allows mounting directories from the host system. This way, an analysis tree can be mounted and the driver scripts can be run from a Docker container. Preserving the Docker image along an analysis tree enables the user to reproduce the analyses using the same software environmnent used to generate it in the first place.

## DISCUSSION

The framework presented here provides a simple and intuitive way to organize computational analyses. The user doesn’t need to learn new syntax to define workflows. This is done instead by laying out data files and scripts in a directory structure, according to simple principles. Maintaining the whole analysis tree under version control makes it easier to experiment with analysis steps and collaborate with other researchers. This framework is in development on github (https://github.com/apuapaquola/rf.git).

## ACKNOWLEGEMENTS

NIH MH095741, NIH MH088485, Helmsley Charitable Trust.

## REFERENCES

Köster J, Rahmann S. Snakemake-a scalable bioinformatics workflow engine. Bioinformatics. 2012 Oct 1;28(19):2520–2. Epub 2012 Aug 20. PubMed PMID: 22908215.

Ebert P, Müller F, Nordström K, Lengauer T, Schulz MH. A general concept for consistent documentation of computational analyses. Database (Oxford). 2015 Jun 8;2015:bav050. doi:10.1093/database/bav050. Print 2015. PubMed PMID: 26055099; PubMed Central PMCID: PMC4460408.

Goecks J, Nekrutenko A, Taylor J; Galaxy Team. Galaxy: a comprehensive approach for supporting accessible, reproducible, and transparent computational research in the life sciences. Genome Biol. 2010;11(8):R86. doi: 10.1186/gb-2010-11-8-r86. Epub 2010 Aug 25. PubMed PMID: 20738864; PubMed Central PMCID: PMC2945788.

Noble WS. A quick guide to organizing computational biology projects. PLoS Comput Biol. 2009 Jul;5(7):e1000424. doi: 10.1371/journal.pcbi.1000424. Epub 2009 Jul 31. PubMed PMID: 19649301; PubMed Central PMCID: PMC2709440.

Goodstadt L. Ruffus: a lightweight Python library for computational pipelines. Bioinformatics. 2010 Nov 1;26(21):2778–9. doi: 10.1093/bioinformatics/btq524. Epub 2010 Sep 16. PubMed PMID: 20847218.

Trapnell C, Roberts A, Goff L, Pertea G, Kim D, Kelley DR, Pimentel H, Salzberg SL, Rinn JL, Pachter L. Differential gene and transcript expression analysis of RNA-seq experiments with TopHat and Cufflinks. Nat Protoc. 2012 Mar 1;7(3):562–78. doi: 10.1038/nprot.2012.016. Erratum in: Nat Protoc. 2014 Oct;9(10):2513. PubMed PMID: 22383036; PubMed Central PMCID: PMC3334321.

